# Positive strand RNA viruses differ in the constraints they place on the folding of their negative strand

**DOI:** 10.1101/2022.02.01.478670

**Authors:** Morgan R. Herod, Joseph Ward, Andrew Tuplin, Mark Harris, Nicola J. Stonehouse, Christopher J. McCormick

**Affiliations:** School of Molecular and Cellular Biology, Faculty of Biological Sciences and Astbury Centre for Structural Molecular Biology, University of Leeds, Leeds, UK; Clinical and Experimental Sciences, Faculty of Medicine, University of Southampton, Sir Henry Wellcome Laboratories, University Hospital Southampton, Southampton SO16 6YD, UK; Institute for Life Sciences, University of Southampton SO17 1BJ, United Kingdom

## Abstract

Genome replication of positive strand RNA viruses requires the production of a complementary negative strand RNA that serves as a template for synthesis of more positive strand progeny. Structural RNA elements are important for genome replication, but while they are readily observed in the positive strand, evidence of their existence in the negative strand is more limited. We hypothesised that this was due to viruses differing in their capacity to allow this latter RNA to adopt structural folds. To investigate this, ribozymes were introduced into the negative strand of different viral constructs; the expectation being that if RNA folding occurred, negative strand cleavage and suppression of replication would be seen. Indeed this was what happened with hepatitis C virus (HCV) and feline calicivirus (FCV) constructs. However, little or no impact was observed for chikungunya virus (CHIKV), human rhinovirus (HRV), hepatitis E virus (HEV) and yellow fever virus (YFV) constructs. Reduced cleavage in the negative strand proved to be due to duplex formation with the positive strand. Interestingly, ribozyme-containing RNAs also remained intact when produced *in vitro* by the HCV polymerase, again due to duplex formation. Overall, our results show that there are important differences in the conformational constraints imposed on the folding of the negative strand between different positive strand RNA viruses.

## INTRODUCTION

Positive strand RNA viruses infect all forms of life and impose significant environmental, economic and social hardship due to the diseases that they cause. For those viruses that infect higher eukaryotic organisms, viral genome replication occurs in membrane rich compartments within the cytosol often referred to as replication complexes (RCs) (1). These compartments are thought to concentrate the viral and host proteins needed for genome replication (1,2) as well as protect the viral RNAs produced in them from intrinsic immune sensors and the action of antiviral proteins (3,4).

The basic principles of viral genome replication were first determined by pulse chase experiments that allowed RNA synthesis to be followed over time (5–11). The current paradigm, based on these studies, is that genomic RNA ([+] RNA) is first recruited to the RC before serving as a template for production of a complementary negative strand RNA ([-] RNA) intermediate. This [-] RNA is closely associated with the [+] RNA from which it was synthesized, forming a double stranded RNA (dsRNA) complex on extraction from cells that is referred to as the replicative form (RF) (6,11). The [-] RNA within the RF subsequently serves as a template for [+] RNA synthesis. Production of [-] and [+] RNA is asymmetric, with multiple copies of the [+] RNA at various stages of synthesis typically being found associated with a single [-] RNA template. In this state the RNA species present is known as a replicative intermediate (RI) and when extracted from cells appears as a double stranded structure with multiple single stranded branches extending from it, with each branch representing a partially synthesized nascent [+] RNA (5,7–10). Although formal demonstration of RF and RI formation has typically been restricted to those viruses that show robust replication, this process is believed to be the same for all positive strand RNA viruses infecting higher eukaryotic organisms, but adapted or extended for those viruses producing subgenomic RNAs (sgRNAs) in addition to their full length genome.

Intramolecular RNA structures encoded within the viral genome play a key role in the viral replication cycle. Many viruses use internal ribosome entry sites (IRESes) as a substitute for a 5’ cap to ensure translation of their viral proteins (12,13). Use of stem loops within coding regions to control frameshifting is also common (14–16). Importantly, RNA structure is an integral part of many *cis*-acting replication elements (CREs); RNA elements that play a more direct role in genome replication itself. The function of these CRE elements range from controlling the switch between translation to replication (17–20), acting as co-factors in enzymatic processes such as viral protein uridylation (21) and serving as promoters to direct initiation of viral polymerase activity (22–25).

The single stranded nature of the viral genome provides the CREs within it the relative freedom to fold. However, once [-] RNA synthesis has occurred, constraints are encountered. This is because the complementary nature of the [-] and [+] RNAs promotes dsRNA formation. Despite this several viruses harbour structured CREs in their [-] RNA; CREs that act as promoters for genomic and subgenomic RNA production (25–29). Thus, at some point the duplex base pairing masking these CREs has to be separated in such a way that intramolecular base pairing is promoted while intermolecular base pairing is prevented. To date, the only way used to monitor RNA folding in the [-] RNA has been to employ CRE-dependent replication as a read-out (25,26,29). As CRE function is specific to the virus, this does not offer a uniform way to examine RNA folding across the [-] RNA of different viruses. Indeed, to our knowledge, the issue of whether sequences other than CRE elements are able to adopt an RNA fold within the [-] strand has never been addressed. Part of the reason is that many RNA structural analysis techniques require disruption of the RC, an action which facilitates the collapse of RF and RI forms into a double stranded state (30,31). Furthermore, the few techniques that allow RNA structure to be examined *in situ* (31,32) are hampered by the typical low abundance of the RF and RI in the infected cell. Thus any technique that offers the ability to monitor RNA folding *in situ*, and to be able to extend this analysis to structures beyond the confines of CRE elements, has the potential to provide valuable insight into the inner workings of the viral RC.

Ribozymes are self-cleaving RNAs whose activity depend on both RNA secondary structure and other higher ordered RNA interactions (33). We reasoned that embedding a ribozyme (Rbz) in the [-] RNA and monitoring the extent to which this strand is subsequently cleaved would provide a direct *in situ* readout of RNA folding. In this study we used Rbzs to assess RNA folding within the [-] RNA of hepatitis C virus (HCV), feline calicivirus (FCV), chikungunya virus (CHIKV), hepatitis E virus (HEV), human rhinovirus (HRV) and yellow fever virus (YFV). Our results confirm that the [-] strand of some positive strand RNA viruses are able to form functional RNA structures. Excitingly, we also demonstrate that this is not conserved across all positive strand RNA viruses but is specific to different divergent families and genera. Finally, our data show that for sequences in the [-] RNA to adopt a structural fold, active participation of host and/or viral proteins is required.

## MATERIAL AND METHODS

### Cell culture

HeLa(Ohio), CRFK and BHK21 cells were maintained in DMEM (Invitrogen) supplemented with 50 units penicillin, 50 mg streptomycin and 10%(v/v) foetal calf serum at 37°C and 5% CO_2_. Huh7.5 cells were maintained under similar conditions but with an additional supplementation of non-essential amino acids made to the media.

### DNA constructs

The JFH-1 based monocistronic replicons, JFH1DVR-mono and JFH1mono(GAA), have been described (34) and served as HCV gt2a replication functional and polymerase defective controls lacking Rbzs. To insert the positive strand hepatitis delta virus (HdV) Rbz sequence into JFH1DVR-mono, PCRs using primer pairs 1 + 2 (see Table S1 for primer sequences) with template pMNV* (35), and primer pairs 3 + 4 and 5 + 6 with template JFH1DVR-mono were used to generate 3 overlapping products that were then combined in a second round PCR reaction with primer pairs 3 and 6. This product and a comparable DNA containing an inactive HdV Rbz, introduced using primer pairs 7 + 8, were cloned into the JFH1DVR-mono containing plasmid via *Bgl*II and *Rsr*II restriction sites to generate plasmid constructs encoding HCVgt2a_HdV(wt) and HCVgt2a_HdV(ko).

To exchange the HdV Rbz for the satellite RNA of tobacco ringspot virus (sTRSV) Rbz (36) and the cleavage optimized *Shistosoma mansoni* N79 Rbz (37), PCRs were performed with primer pairs 9 + 10 and 11 + 12 using HCVgt2a_HdV(wt) as a template. The two DNA products generated were used as templates alongside Ultramer oligonucleotides TRSV or N79 (Integrated DNA Technologies) in a second round of PCR employing primer pairs 9 + 12. Resultant products were cloned directly into the HCVgt2a_HdV(wt) plasmid via *Bgl*II and *Rsr*II restriction sites to generate HCVgt2a_sTRSV(wt) and HCVgt2a_N79(wt) containing plasmids. The same products were also further modified using another round of 2-step PCR mutagenesis using internal primer pairs 13 + 14 or 15 + 16 alongside 9 and 12 before again cloning into the HCVgt2a_HdV(wt) plasmid to generate HCVgt2a_sTRSV(ko) and HCVgt2a_N79(ko) containing plasmids.

To add a second copy of the N79 Rbz between the NS5A and NS5B coding region of the HCVgt2a replicon, three first round PCRs were set up using primer pairs 17 + 18, 19 + 20 and 21 + 22. The template used in each reaction was either HCVgt2a_N79(wt) or HCVgt2a_N79(ko). All three products were combined in a second round PCR using primer pairs 17 + 22 and the resultant DNA cloned into both the HCVgt2a_N79(wt) or HCVgt2a_N79(ko) containing plasmids via *Sac*II and *Sal*I restriction sites to generate plasmids containing HCVgt2a_2xN79(wt/wt), HCVgt2a_2xN79(ko/ko), HCVgt2a_2xN79(wt/ko) and HCVgt2a_2xN79(ko/wt).

Insertion of the N79 Rbz into other replicon constructs was achieved as follows. For the HRV replicons, PCRs were undertaken using primer pairs 23 + 24 in combination with either HCVgt2a_N79(wt) or HCVgt2a_N79(ko) as a template, as well as primer pairs 25 + 26 in combination with pR16.11 (Accession number L24917.1; a gift from T. Tuthill) as a template. The resultant sets of DNAs were combined in a second round PCR with primers 23 + 26 and the resultant product cloned into an in-house pR16.11 derived replicon (sequence available on request) via *Asc*I and *Cla*I restriction sites to generate plasmids containing HRV_N79(wt) and HRV_N79(ko). A control replication defective version of HRV_N79(ko) was generated by cutting the associated vector with *Nsi*I, polishing with Phusion polymerase (NEB) and religating. For the YFV replicons, PCRs were undertaken using primer pairs 27 + 28 and 29 + 30 using pACNR-FLYF-17D (38) (a gift from P. Bredenbeek) as a template as well as primer pairs 31 + 32 using either HCVgt2a_N79(wt) or HCVgt2a_N79(ko) as a template. The resultant 3 sets of DNAs were combined in a subsequent PCR with primer pairs 27 + 30 and the resultant product cloned into pACNR-FLYF-17D via *Not*I and *Mlu*I restriction sites to generate plasmids encoding for the YFV_N79(wt) and YFV_N79(ko) replicons. A control replication defective version of YFV_N79(ko) was generated by cutting the vector with *Cla*I, polishing with Phusion polymerase and religating. For the CHIKV replicons, PCR products was generated using primer pairs 33 and 34 in combination with either YFV_N79(wt) or YFV_N79(ko) as templates. The resultant product was cloned into pSP6_ChikRepI-PRlucSG-FlucWT or pSP6_ChikRepI-PRlucSG-FlucGAA (39) via *Apa*I + *Bln*I to produce plasmids containing CHIKV_N79(wt), CHIKV_N79(ko) and CHIKV_N79(ko)GAA replicons.

For the HEV replicons the GFP coding region from pSK-E2-GFP (40)(a gift from P. Farci) was first replaced by nano-luciferase to generate a nano-luciferase expressing replicon. To achieve this, PCR was used to amplify the nano-luciferase open reading frame (ORF) using primers 35 + 36. The upstream HEV sequence was amplified by PCR using 37 + 38. The resulting PCR fragment were combined in a second round PCR and cloned using *Xho*l and *EcoR*I to generate pSK-E2-nLuc. To insert the Rbz into pSK-E2-nLuc, PCRs were undertaken using primer pairs 39 + 40 using either HCVgt2a_N79(wt) or HCVgt2a_N79(ko) as a template. The resultant product was cloned into pSK-E2-nLuc via *EcoR*I restriction sites to generate plasmids containing the replicons HEV-nLucN79(wt) and HEV-nLucN79(ko). A control replication defective version of pSK-E2-nLuc containing an inactivating mutation to the RNA-polymerase active site (GDD>GNN) was generated by PCR using primer 41 + 42 and 43 + 44 with pSK-E2-nLuc as template. The resultant PCR products were combined in a second round PCR using primers 41 + 44 and cloned using *Xho*I and *EcoR*I to generate a plasmid containing the replicon HEV-nLuc-GNN.

For the HCV genotype 1b replicon, first round PCR products were generated using primer pairs 45 + 46 with template HCVgt2a_N79(wt) or HCVgt2a_N79(ko), and using primer pairs 47 + 48 with template pFKI341PVIlucUbiNS3-3’dgET (41) (a gift from R. Bartenschlager). Resultant DNAs were used a templates in a second round PCR using primer pairs 45 + 48 and the products produced cloned into pFKI341PVIlucUbiNS3-3’dgET via *Not*I and *BssH*I restriction sites to generate plasmids encoding HCVgt1b_N79(wt) and HCVgt1b_N79(ko) replicons.

For the FCV constructs, pQ14 (42) (a gift from K. Green) was used as a template in a 2-step PCR using primer pairs 49 + 50 and 51 + 52. The first round products were pooled and used as a template with the primer pairs 49 + 52 to generate a DNA that was cloned back into pQ14 via *BstB*I and *Spe*I restriction sites, introducing a multiple cloning site (MCS) into the leader sequence of FCV. GBlocks (Integrated DNA Technologies) encoding a Nanoluc_antisense_sTRSV fusion product or Nanoluc_antisense_N79 fusion product were then cloned into this MCS via *Kpn*I and *Pst*I restriction sites generating plasmids containing the viruses FCV_sTRSV(wt) and FCV_N79(wt). The GBlocks were also used as templates for a two-step PCR using internal mutagenic primers 13 and 14 or 15 and 16 to generate products that were cloned into the MCS, generating FCV_sTRSV(ko) and pQ14_FCV_N79(ko) containing plasmids. A replication-defective FCV construct was generated by cutting pQ14 with *Xho*I, polishing with Phusion polymerase and religating.

To modify plasmids containing the N79-containing replicon constructs so that they could be used to transcribe the [-] strand, these DNAs were first linearised with the restriction site found downstream and adjacent to their 3’UTR (*Not*I, *Xba*I, *Bgl*II, *BamH*I, and *Xho*I for CHIKV, HCVgt2a, HEV, HRV and YFV constructs respectively). A pair of complementary oligos containing either a SP6 (HCV, HEV, HRV and YFV) or T7 (CHIKV) promoter and with overhanging ends compatible with those of the digested plasmid DNA were ligated into these same DNAs. The presence and orientation of the inserted DNA was confirmed by PCR and sequencing. For the purpose of identification, the names of the constructs produced bear the parental constructs name from which they derive but contain an additional “as” (antisense) prefix (e.g. asHCVgt2a-N79(wt)).

To generate constructs containing a reverse complemented N79 sequence to be used for transcribing template RNAs for *in vitro* NS5B RdRp assays, a 2-step PCR was performed. Primer pairs 53 + 54 and 55 + 56 generated amplified product from HCVgt2a_N79(wt) and HCVgt2a_N79(ko) templates. These 2 reaction products were then combined in a second PCR reaction with primer pairs 53 + 56 and the resultant DNA cloned into pCR-Blunt (Invitrogen) before being excised using *Xba*I + *EcoR*I and cloned into *Xba*I + *EcoR*I cut pSGRJFH1luc, generating pT7HCV(3’-)341asN79^wt^ and pT7HCV(3’-)341asN79^ko^. Constructs were linearized with *Mfe*I and polished with Mung Bean nuclease prior to use in T7 transcription reactions that generated RNA template for subsequent NS5B RNA-dependent RNA polymerase (RdRp) reactions.

To generate control constructs for producing the same RNA species to be synthesized in the NS5B RdRp reactions, but via T7 polymerase, PCRs were performed using the primer pair 57 + 54 in combination with pSGRJFH1luc as template DNA, and the primer pair 55 + 58 in combination with either HCVgt2a_N79(wt) or HCVgt2a_N79(ko) as template DNAs. First round products were combined in a second PCR using primers 57 and 58. Products were first cloned into pCR-Blunt then excised with *Xba*I and *EcoR*I and cloned into *Xba*I + *EcoR*I cut pSGRJFH1luc to produce pT7HCV(5’+)341N79^wt^ and pT7HCV(5’+)341N79^ko^.

### Production of RNA transcripts from DNA templates

All RNAs other than HEV and FCV transcripts were generated as follows. Five micrograms of linearised plasmid template were pre-treated with RNAsecure (ThermoFisher) and used in a 50μl transcription reaction containing 5 x transcription buffer (ThermoFisher), 8mM rNTPs, 1.25 units RiboLock RNAse inhibitor (ThermoFisher), and 60 units of either T7 or SP6 RNA polymerase (ThermoFisher). After an overnight incubation at 30°C, 2 units RNA-free DNAse (NEB) was added and reactions left at 37°C for 30 minutes. RNA transcripts were precipitated by addition of LiCl and the resultant pellet washed in 70% ethanol before resuspension in RNase-free water. Capping of YFV and CHIKV replicon RNA transcripts was performed using the Vaccinia Capping System (NEB) according to manufacturer’s recommendations.

HEV and FCV transcripts were generated using the HiScribe T7 ARCA mRNA kit (NEB) using 1 μg of linearised plasmid template following manufacturer’s instructions. The integrity of all RNAs was verified by running RNAs on a MOPS-formaldehyde agarose gel and visualizing product by SybrGold staining (ThermoFisher). When required, relative quantification of RNA species on the gels was achieved using a ChemiDoc XRS+ imager (Biorad).

### Cell electroporation and monitoring of replicon replication

Huh7.5 cells were used to monitor replication of HCV, HEV and YFV replicons, BHK21 used to monitor replication of CHIKV replicons, HeLa(Ohio) used to monitor replication of HRV replicons and CRFK cells used to monitor replication of FCV. Once cells had reached confluence they were detached, washed x 2 in ice-cold RNase-free phosphate buffered saline (PBS) and resuspended at 1 x 10^7^ cells /ml in ice-cold PBS. Four hundred microliters of the cell suspension was mixed with 2 μg replicon transcript, transferred to a 0.4cm cuvette, electroporated at 270V, 960μF (Biorad GenePulse II) and resuspended in 6ml growth medium. Cells were plated out in 12-well plates and maintained at 37°C (HCV, YFV, CHIKV, HEV) or at 33°C (HRV) until harvested by washing in PBS and lysis in Passive Lysis Buffer (Promega). Luciferase activity was determined using either the Renilla Luciferase Assay Kit (Biotium), Nano-Glo Luciferase Assay System (Promega) or Dual Luciferase Reporter Assay (Promega). Live cell imaging was performed with an IncuCyte Dual Colour Zoom FLR (Essen BioScience) within a 37□°C humidified CO2 incubator scanning hourly up to 120 hr post-transfection, collecting multiple images per well. The .mp4 files generated had individual stills extracted from them using VLC media player.

### Northern Blot

To generate probes, the HCVgt2a_HdV(wt) vector was used a template in an asymmetric PCR reaction (43) containing a pair of outer primers (primers 59 and 60) and either an inner forward (primer 61) or inner reverse primer (primer 62). Single stranded products produced by the reaction were separated from other reaction products on an agarose gel and purified. Resultant DNAs as well as a 316bp GAPDH PCR product amplified from p-Tri-GAPDH (ThermoFisher) were biotinylated using PlatinumBrightBIO (Kreatech) reagent. Total cellular RNA was recovered using TriFAST reagent (Peqlabs) according to manufacturer’s instructions, run on a 0.8% MOPS-formaldehyde agarose gel, the gel stained using SybrGold to confirm rRNA integrity and the RNAs subsequently transferred to SensiBlot Plus nylon membrane (Fermentas). After uv-crosslinking, membranes were pre-blocked by a 30 minute incubation in Ultrahyb (ThermoFisher) at 42°C, before an overnight incubation in Ultrahyb containing biotinylated probe. Unbound probe was removed by washing the membrane at 42°C in 2xSSPE + 0.1% SDS and subsequently in 0.1XSSPE 0.1% SDS. Bound probe was detected using the BrightStar Northern Blot detection kit (Invitrogen). Because probes could not be stripped from membranes, parallel blots were probed with GAPDH as a control. Images were captured on film. The specificity of both strand specific probes was confirmed by hybridization to control Northern blot membranes containing either HCV [+] or [-] strand *in vitro* RNA transcripts (Fig S1).

### RdRp assay

Five hundred nanograms of HCV(3’-)341asN79^wt^ and HCV(3’-)341asN79^ko^ RNA transcripts were heated to 90°C for 2 minutes in a final volume of 16.2 μl before slowly cooling to room temperature and placing on ice. To these RNA templates the following was added; 2.5μl 10 x NS5B transcription buffer (200 mM Tris-HCl (pH 7.5), 50 mM MgCl_2_, 10 mM DTT, 250 mM KCl, 1 mM EDTA), 0.6 μl RNaseOUT (ThermoFisher), 1μl 10mM rATP, 1μl 10mM rCTP, 1μl 10mM rGTP, 1μl 0.1mM rUTP, 1μl [α-32P]rUTP (10 μCi; 3000Ci/mmol Perkin Elmer) and 0.7μl JFH1 NS5BΔC21 or JFH1 NS5BΔC21(GAA) at 0.3mg/ml (44). Parallel T7 reactions performed at the same time used 1.25 μg *Xba*I linearised pT7HCV(5’+)341N79^wt^ and pT7HCV(5’+)341N79^ko^ as templates in a 25μl final reaction volume containing 2.5μl 10 x T7 transcription buffer (NEB), 0.6μl RNaseOUT, 1μl 10mM rATP, 1μl 10mM rGTP, 1μl 10mM rCTP, 1μl 1mM rUTP, 0.5μl [α-32P]rUTP and 1μl T7 polymerase (NEB). After incubation at room temperature for 2 hours the NS5B RdRp reactions were placed on ice while the T7 reactions were supplemented with 1 unit RNase-free DNase and incubated at 37°C for 20 minutes. All samples were centrifuged through Microspin G25 columns (Amersham) pre-equilibrated with 10mM Tris (pH7.5) 1mM EDTA (TE) buffer before further addition of TE buffer to a final volume of 100μl, phenol/chloroform extraction and ethanol precipitation in the presence of 20μg glycogen. Half of each RNA was kept back for subsequent analysis while the other half was subjected to digestion with RNase A in a 100μl high salt buffer (40 μg/ml RNase A, 10mM Tris (pH7.5), 350mM NaCl, 5mM EDTA) for 60 minutes at 30°C before addition of 4μl 20 mg/ml proteinase K and 6.6μl 10%SDS and incubating at 37°C for 30 minutes. Reaction products were then phenol/chloroform extracted, ethanol precipitated, resuspended in 8μl H_2_O and again half the recovered material was kept back for subsequent analysis. The remaining 4μl of samples were mixed with 4 μl of formamide and the RNAs heated to 85°C for 2 minutes before being rapidly diluted by the addition of 196μl Rbz cleavage buffer preheated to 85°C. Samples were allowed to cool slowly to room temperature before being subjected to ethanol precipitation in the presence of 20μg glycogen as a carrier. RNAs from both treated and untreated arms of the experiment were separated on a 5% denaturing acrylamide:bisacrylamide (19:1) gel alongside a RiboRuler Low Range RNA Ladder (ThermoFisher) and the gel fixed, stained with Methylene Blue and dried onto Whatman 3MM filter paper. Gel images were captured by exposure to both CL-Xposure X-ray film (ThermoFisher) or BAS-MP IP phosphoimager plate (FujiFilm) and scanned at 50 μm resolution at 635 nm in a FujiFilm FLA-5100 fluorescent imager analyser.

## RESULTS

### Cleavage of an HCV replication intermediate by a cis-acting hepatitis delta virus (HdV) Rbz

To assess the potential for Rbzs to fold and cleave the [-] RNA of a positive strand RNA virus *in cis*, we initially chose to focus on HCV. The basis for this decision was because of evidence that HCV allows folding of relatively complex native structures in its [-] RNA (26,29). Additionally we opted to use subgenomic replicon-based constructs because they enable the relevant stages of virus genome replication to be studied in isolation from other stages of the virus replicative cycle - such as entry, packaging and egress. A monocistronic HCV genotype 2a (gt2a) replicon (JFH1DVR-mono) was selected that expressed both a Renilla-FMDV2A luciferase reporter fusion protein and the HCV viral replicase (NS3-5B) needed for RNA replication. It was subsequently modified so as to introduce a reverse complemented HdV Rbz sequence between the Renilla and FMDV2A coding regions, generating HCVgt2a_HdV(wt) (Fig 1a). A mutated HdV Rbz, with nucleotide substitutions designed to disrupt the folding of this RNA and hence its activity, was introduced into the antigenomic strand of an otherwise identical replicon to generate a second control construct, HCVgt2a_HdV(ko). Both constructs had the reverse complemented HdV sequence positioned within their coding region in such a way that the ORF was maintained and the foreign peptide encoded by the insertions was identical. Cells were transfected with these two constructs as well as the original JFH1DVR-mono construct and a polymerase-defective control construct, and luciferase activity was measured over time as a readout of replication activity (Fig 1b).

**Figure 1.**
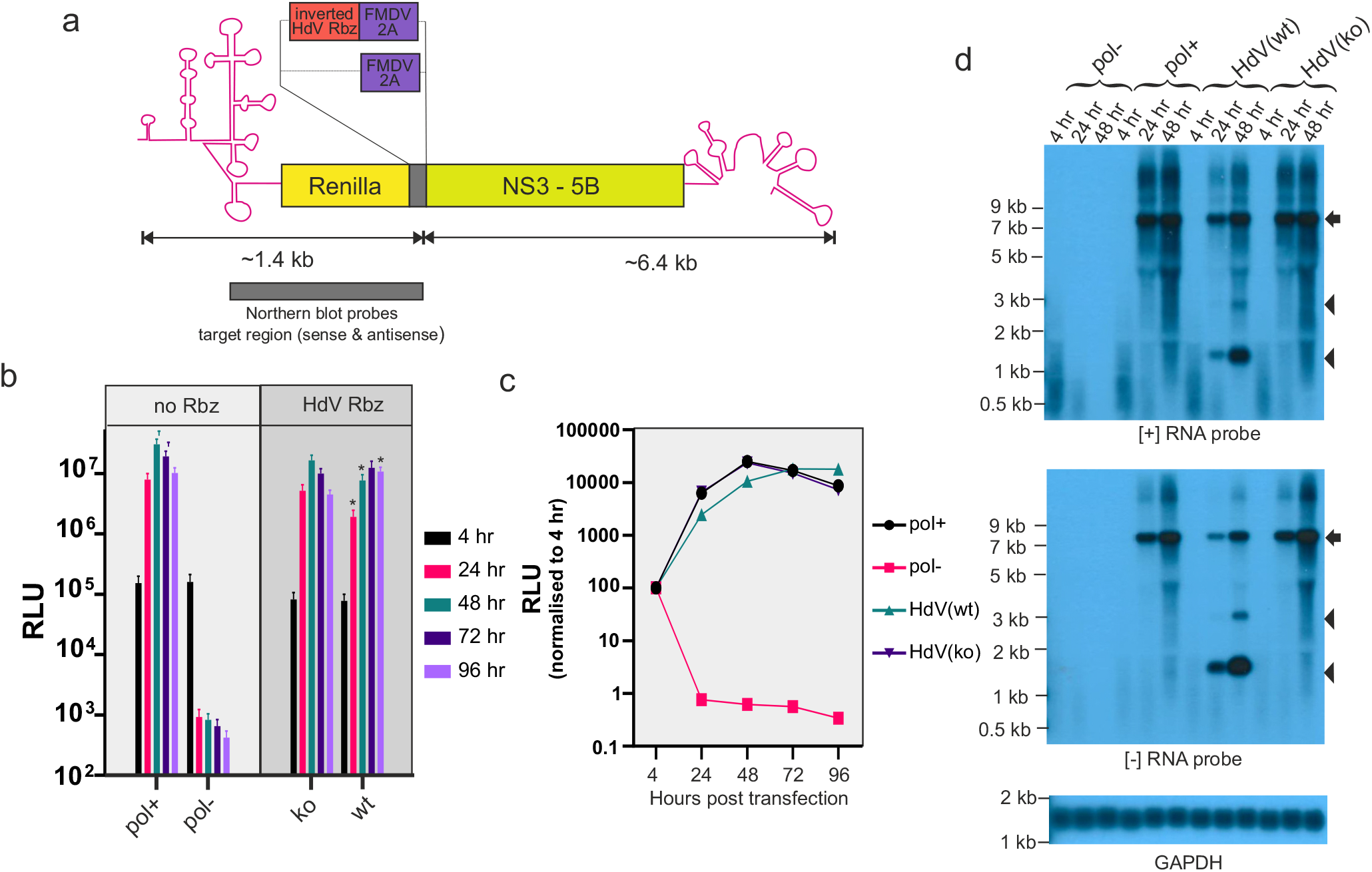
Placing the HdV Rbz in the [-] RNA of HCV cleaves this RNA and suppresses replication. (A) Schematic depiction of the HCV gt2a replicon encoding the HdV Rbz in its [-] RNA. Image includes the region of both [-] and [+] strands recognised by the strand specific probes used for Northern blotting. (B and C) Replication of constructs carrying either an active (wt) or inactive (ko) HdV Rbz. Included are replication competent and replication defective controls lacking the inserted Rbz sequence. Data presented in both graphs are from the same experiments but in (C) they have been normalised to the 4 hr signal. Significant differences between HdV(ko) and (wt) constructs are highlighted in (B) (* p<0.05; paired t-test; n=6). (D) Northern blot of RNA from transfected cells. The arrow highlights the position of full length transcripts and the arrow heads the position of products produced as a result of Rbz activity.

Four hours post-transfection (hpt), when the luciferase activity arises from translation of the input RNAs alone, the signal was 2-fold lower for both the HdV constructs compared to JFH1DVR-mono and replication defective control constructs. Given that no replication would have occurred at this point, as illustrated by the signal being the same between the replication competent and defective controls, this difference likely comes from partial disruption of luciferase activity arising from the insertion of the reverse complemented HdV Rbz sequence into the Renilla-FMDV2A fusion protein. After this 4 hour time point, HCVgt2a_HdV(wt), HCVgt2a_HdV(ko) and JFH1DVR-mono all produced an initial increase in luciferase over time compared to a decrease seen for the polymerase defective control construct, demonstrating that all 3 former RNAs were replication-competent. However, the luciferase signal from the replicon with the active Rbz significantly diverged from its inactive Rbz counterpart as the later produced a more rapid rise in luciferase activity, with luciferase levels peaking at 48 vs. 96 hpt. When relative light unit (RLU) values were normalized to the signal at 4 hours to correct for inherent differences in encoded enzyme activity as well as transfection efficiencies, HCVgt2a_HdV(ko) showed identical replication compared to the parental JFH1DVR-mono construct. As seen with the non-normalized data, replication of HCVgt2a_HdV(wt) was suppressed (Fig 1c), with luciferase levels at 32% and 44% of those produced by its HdV(ko) counterpart at 24 and 48 hpt.

If Rbz activity was suppressing replication then cleavage of the [-] RNA would be anticipated. To establish whether this was the case, RNA was recovered from these same experiments and assessed by Northern blot (Fig 1c). Strand specific probes directed at the Renilla coding region were unable to detect the input RNA at 4 hpt. However, full length [+] and [-] RNA could be detected at 24 and 48 hpt for all replication competent constructs but not the replication defective control. Consistent with the luciferase replication data, the abundance of these two RNAs were somewhat reduced in the HCVgt2a_HdV(wt) compared to HCVgt2a_HdV(ko) and JFH1DVR-mono transfected cells. The appearance of detectable levels of replicon transcript in HCVgt2a_HdV(wt) transfected cells at 24 hours onwards also coincided with the appearance of two other smaller RNA products that were absent from all other experimental groups. Importantly, the size of the most prominent of these matched the size of the [-] RNA fragment that was expected should Rbz cleavage be occurring. Interestingly, a similar size band also appeared in the genomic blot, indicating that the cleaved [-] RNA might serve as a template for RdRp activity. The identity of the second minor RNA species is not clear and interestingly it was not always observed. However, given that it was approximately twice the size of the cleaved 3’ end of the [-] RNA, and was picked up by both probes, it would be consistent with the predicted size of a copy-back product.

### Improved cleavage of the [-] RNA strand using hammerhead Rbzs

Having established by cleavage of the HCV [-] RNA using a *cis*-acting HdV Rbz that functional RNA structure was able to form in the virus antigenomic strand, we were interested in whether other smaller Rbzs with complex secondary structure could be used to further enhance cleavage efficiency. The monocistronic HCV gt2a replicon, JFH1DVR-mono, was again adapted so as to contain one of two different reverse complemented hammerhead Rbzs placed between the Renilla and FMDV 2A coding regions. As before, the ORF of the replicon was maintained and control constructs encoding inactive versions of the same Rbz sequence in their [-] RNA were generated by introduction of synonymous mutations. The two Rbz sequences selected for analysis were from satellite RNA Tobacco Ringspot Virus (sTRSV) (36) and another derived from *S. mansoni* that had been further engineered for enhanced cleavage activity (N79) (37). The four constructs generated were HCVgt2a_sTRSV(wt), HCVgt2a_sTRSV(ko), HCVgt2a_N79(wt) and HCVgt2a_N79(ko). Based on luciferase measurements, replication of constructs carrying the active versions of these Rbzs was significantly suppressed (Fig 2a, b), more so than constructs bearing the HdV Rbz. In contrast, the inactive Rbz control constructs demonstrated robust replication, which after normalizing to the input signal at 4 hours matched that of JFH1DVR-mono. Normalised Renilla values produced from HCVgt2a_sTRSV(wt) were 5%, 20% and 78% of those produced from HCVgt2a_sTRSV(ko) at 24, 48 and 72 hours (Fig 2c). More marked was the suppression seen with the N79(wt) Rbz where luciferase activity was found to be 0.1%, 0.2% and 0.4% of the N79(ko) control levels at 24, 48 and 72 hours respectively. Northern blot analyses of RNAs taken from cells 48 hours post transfection and hybridized to probes complementary to the Renilla encoding region of the [+] and [-] strands identified sTRSV Rbz cleaved products (Fig 2d). The relative abundance of these products compared to their full length uncleaved [+] and [-] RNA counterparts was notably increased compared to that seen with the HdV Rbz, confirming that the sTRSV Rbz was indeed more efficient at cleaving the [-] strand of HCV. Also consistent with the luciferase data, the overall levels of transcript were much lower in cells transfected with HCVgt2a_sTRSV(wt) vs. HCVgt2a_sTRSV(ko). No replicon RNA signal was detected in cells transfected with HCVgt2a_N79(wt). Given that replication of HCVgt2a_N79(ko) was robust, this is presumably because of efficient N79 Rbz cleavage suppressing replication levels to below that detectable by Northern blot.

**Fig 2.**
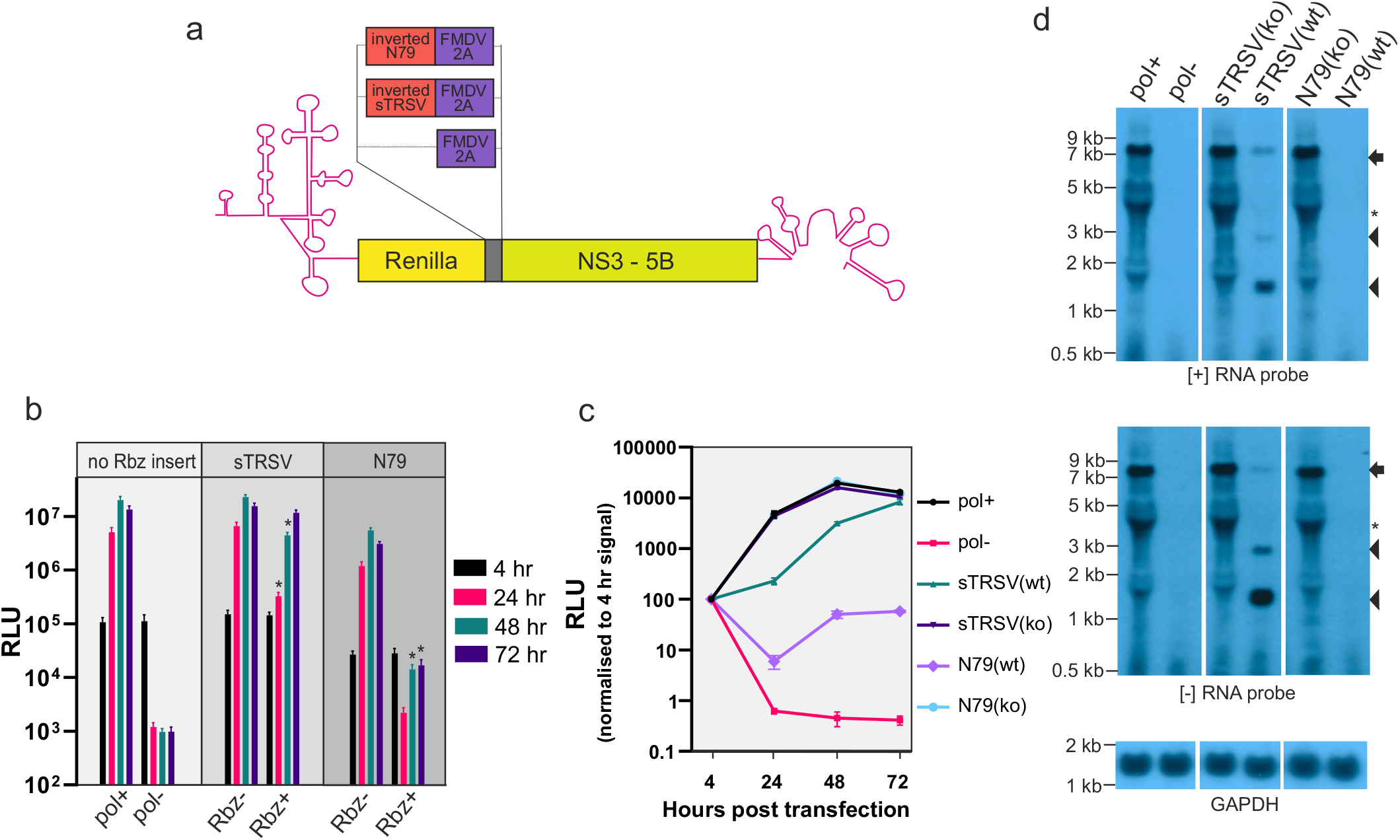
Hammerhead Rbzs are highly effective at cleaving the [-] RNA of HCV replicon constructs. (A) Schematic depiction of the HCV gt2a replicons used in this experiment. (B and C) Replication of constructs carrying either an active (wt) or inactive (ko) sTRSV or N79 hammerhead Rbz. Included are replication competent and replication defective controls lacking the inserted Rbz sequence. Data presented in both graphs are from the same experiments but in (C) they have been normalised to the 4 hr signal. Significant differences between sTRSV(ko) and (wt) constructs, and between N79(ko) and (wt) constructs, are highlighted in (B) (* p<0.05; paired t-test; n=3). (D) Northern blot of RNA from cells transfected with replicon constructs 48 hours earlier. The arrow highlights the position of full length transcripts and the arrow heads the position of products produced as a result of Rbz activity. Other bands on the gel (*) coinicident with the position of ribosomal RNAs, represent background artefacts.

### Rbz mediated suppression of HCV replication is positionally independent

In order to confirm the availability of the [-] RNA strand to form functional RNA structure in other regions of the virus genome during HCV replication, the positional independence of the Rbz was investigated. To achieve this, a second reverse complemented N79(wt) or N79(ko) sequence was introduced into the NS5A/5B boundary of both HCVgt2a_N79(wt) and HCVgt2a_N79(ko) (Fig 3a). This was done in such a way that the NS3-5B replicase would still be expressed and the peptide resulting from this insertion would be cleaved away from both NS5A and NS5B. The resulting four constructs, HCVgt2a_2xN79(wt/wt), HCVgt2a_2xN79(ko/ko), HCVgt2a_2xN79(wt/ko) and HCVgt2a_2xN79(ko/wt), were then transfected into cells and luciferase activity monitored. Consistent with the N79 Rbz being active at both sites, transfection with either HCVgt2a_2xN79(wt/ko) or HCVgt2a_2xN79(ko/wt) resulted in a profound drop in luciferase activity over time, with this being more pronounced in the latter construct such that luciferase values were comparable to those of the replication-defective control (Fig 3b). HCVgt2a_2xN79(wt/wt) also produced comparable luciferase values to the negative control where as HCVgt2a_2xN79(ko/ko) showed increasing luciferase activity for the first 48 hours of the assay that was significantly raised above that of the other 2xN79 constructs. Normalizing the signals to the 4 hour time points revealed that HCVgt2a_2xN79(ko/ko) had slightly reduced replication capacity to that of JFH1DVR-mono, likely resulting from the inserted sequence placed between the duplicated NS5A/5B boundary (Fig 3c). However, given it was still clearly replication competent we conclude that the lack or near lack of replication seen with HCVgt2a_2xN79(wt/ko), HCVgt2a_2xN79(ko/wt) and HCVgt2a_2xN79(wt/wt) is a direct consequence of N79 Rbz activity.

**Fig 3.**
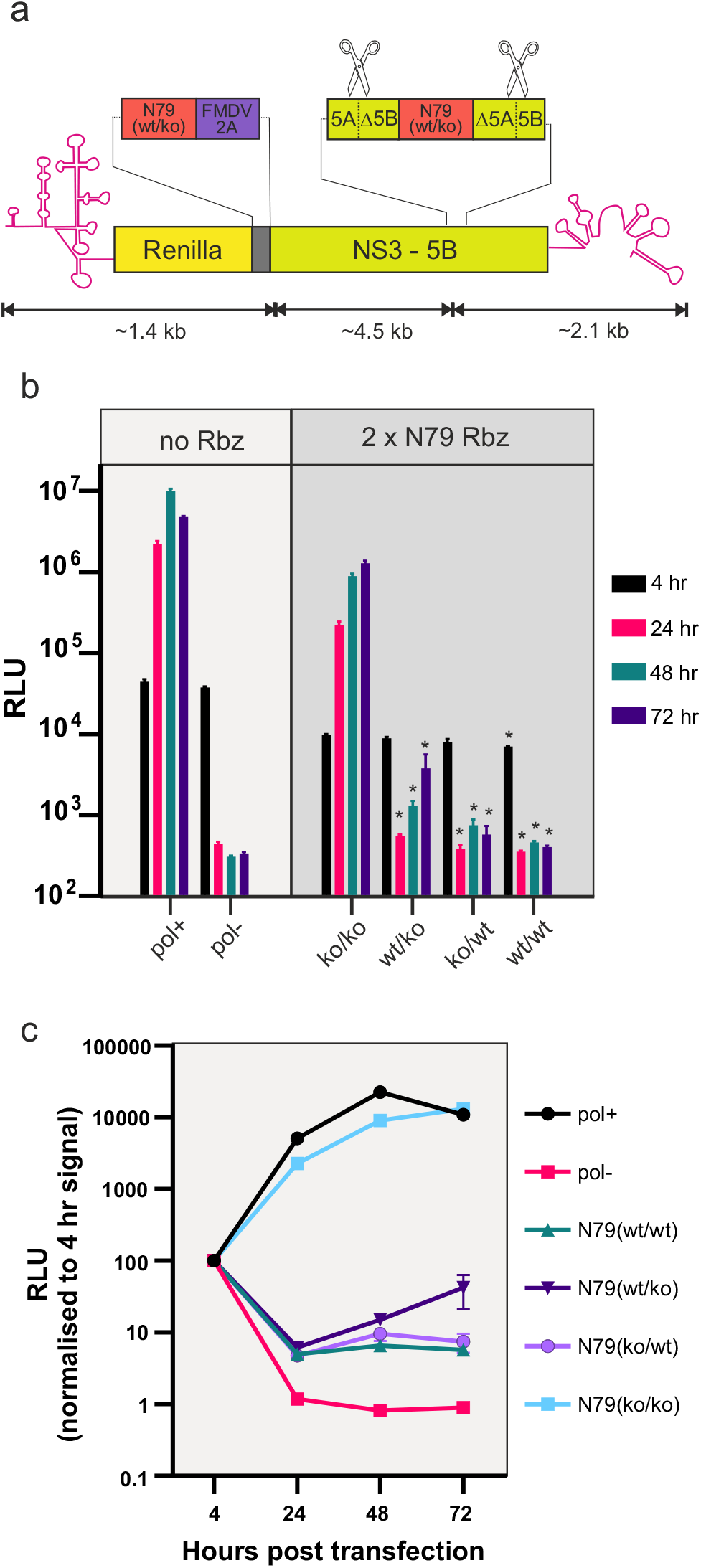
Rbzs embedded in the [-] RNA of HCV suppress replication irrespective of the position they are located. (A) Schematic depiction of HCV gt2a replicons carrying two copies of the N79 Rbz positioned at two different locations within the genome. (B and C) Replication data from the constructs depicted in (A) as well as from replication competent and replication defective controls lacking the inserted Rbz sequence. Data presented in both graphs are from the same experiments but in (C) they have been normalised to the 4 hr signal. Significant differences between the (ko/ko) construct and other 2 x N79 containing constructs are highlighted in (B) (* p<0.05; one-way ANOVA; n=3). No significant differences were observed between other 2 x N79 experimental groups.

### Assessing the impact of the N79 Rbz on the replication of other positive strand RNA virus constructs

Having established that the N79 Rbz effectively suppressed replication of the HCV gt2a replicon, we assessed its impact on replication when placed in the [-] RNA of other positive strand RNA virus constructs. A variety of HCV genotype replicons exist although they display reduced levels of replication compared to JFH-1 gt2a constructs. Introduction of a functional N79 reverse complemented sequence into a HCV gt1b replicon abolished replication whereas an inactive N79 Rbz did not (Fig 4a).

**Fig 4.**
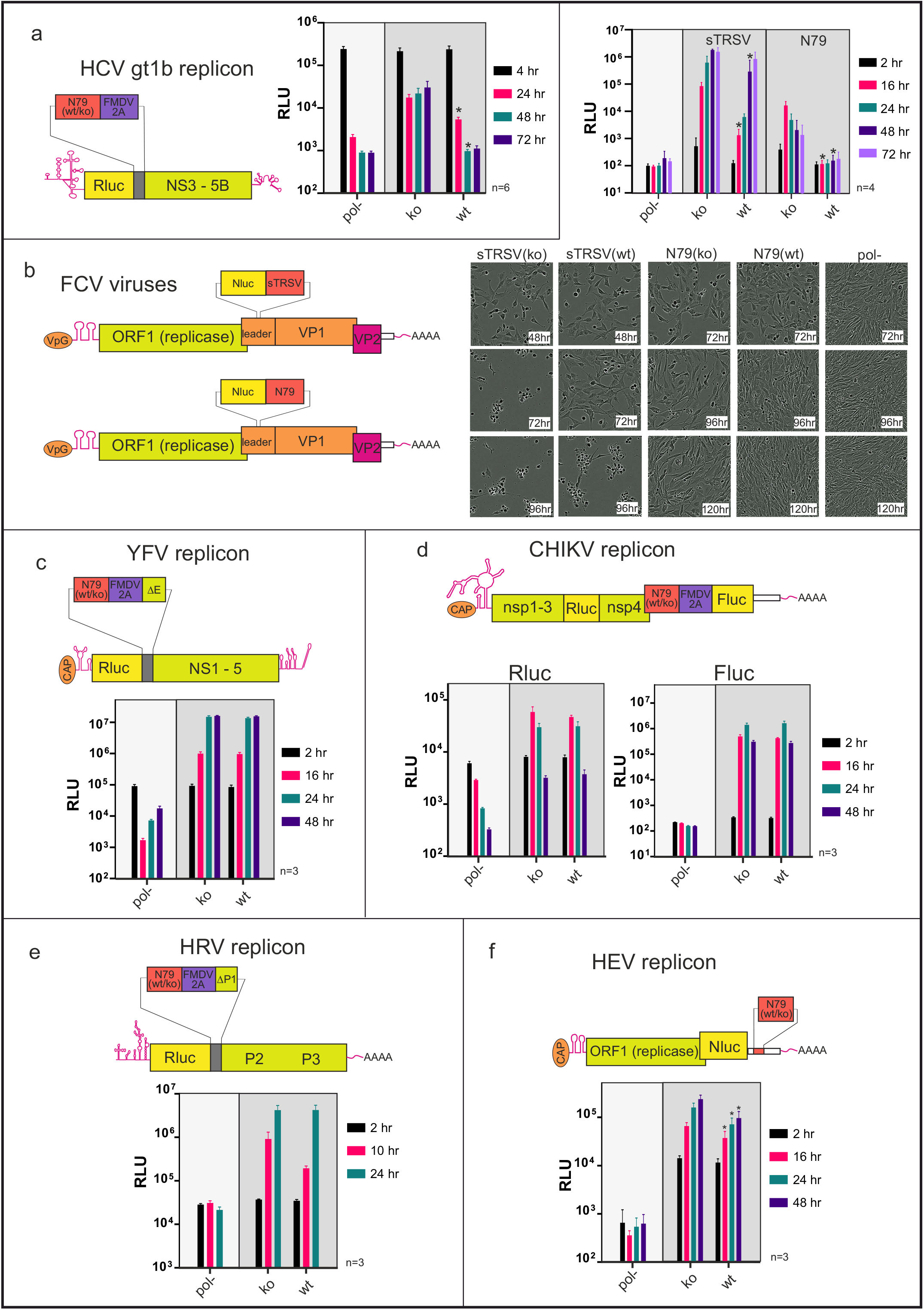
Embedding Rbzs in the [-] RNA of a diverse set of positive strand RNA virus constructs results in different replicative outcomes. A schematic depiction showing the positioning of the N79 Rbz or the sTRSV Rbz is provided for an (A) HCV gt1b replicon (B) FCV genome (C) YFV replicon (D) CHIKV replicon (E) HRV replicon and (F) HEV replicon. Also shown are luciferase replication data from those same isolates and in the case of the FCV virus experiments, representative images illustrating differences in cpe development over time. The number of experimental repeats for each luciferase assay is indicated next to the respective graph. Significant differences between (ko) and (wt) constructs are highlighted (* p<0.05; paired t-test).

We next examined Rbz activity in the [-] RNA of a member of the *Caliciviridae* family; viruses which harbour relatively large stem loops in this strand that function as a subgenomic promoter (45). Because *Caliciviridae* replicons have limited replicative capacity, reverse complemented sTRSV and N79 sequences were instead introduced into the leader sequence of an infectious clone of FCV, a location previously shown to tolerate foreign sequence (46). A Nanoluc coding region was also introduced at this location to simplify monitoring of replication. Transfection of these constructs into cells produced different results depending on the Rbz sequence used (Fig 4b). For constructs carrying either an active or inactive sTRSV Rbz, transfected cells exhibited cytopathic effects (cpe), suggestive of infectious virus production. Importantly, for sTRSV(ko) transfected cells the appearance of cpe always occurred sooner than in sTRSV(wt) transfected cells, although the exact timings of when it was seen did vary between experiments. This earlier appearance in cpe seen for the sTRSV(ko) construct paralleled what was observed for the luciferase activities, with this same construct producing significantly higher levels than sTRSV(wt) at all time points up to and including 48 hours. In contrast, FCV constructs carrying the N79 Rbz sequences in their [-] RNA failed to produce cpe, although there was a suggestion that cell growth was perhaps suppressed, more so for the N79(ko) construct. Monitoring luciferase activity demonstrated that genome replication was occurring, at least for the N79(ko) construct, as luciferase levels increased over the first 16 hours to ~100-fold above background levels and then slowly decreased. In contrast, luciferase activity of the N79(wt) and the replication control construct were similar, demonstrating that the active N79 Rbz predominately blocked replication; a result comparable to what had been observed in the HCV constructs.

Having established that the reverse completed N79 Rbz effectively blocked replication of HCV and FCV, we introduced it into other positive strand virus replicons in order to investigate the conservation of [-] RNAs ability to form functional RNA structures across divergent viruses. Interestingly, for a replicon derived from the 17D vaccine strain of YFV, no difference in replication could be discerned between constructs carrying the functional vs non-functional N79 sequence (Fig 4c). A similar observation was made with the CHIKV replicon, where the Firefly luciferase signal (indicative of subgenomic RNA production) and the Renilla signal (indicative of genomic RNA transcription) were the same for both the functional and inactive N79 constructs at all time-points tested (Fig 4d). The situation was slightly different for HRV, where introduction of a functional N79 Rbz caused a consistent drop (55-88%) in luciferase activity at 12 hours in cells transfected compared to an inactive N79 control construct (Fig 4e). However, this difference between the 2 constructs did not reach significance and disappeared at the 24 hour time point when luciferase activities had peaked. In contrast, replication of the HEV replicon carrying the functional N79 Rbz was significantly impaired, generating a luciferase signal at 24, 48, 72 and 96 hours post transfection that was 92%, 60%, 53% and 44% of the signal produced from the inactive N79 control (Fig 4f). Notably, these experiments showed that irrespective of whether an active N79 Rbz suppressed replication in these four viruses, the reduction in replication, if any, was slight in comparison to either FCV or HCV where N79 imposed a complete or near-complete block on replication.

Rbz folding and hence activity can be influenced by adjacent sequences. Care had been taken to minimize the chance of surrounding sequences interfering with N79 activity by placing ‘insulator’ sequences either side of the Rbz boundaries. However, it remained possible that mis-folding could account for some of the differences observed between the viral constructs regarding N79 activity. To investigate this, we first looked at folding using *in silico* free energy minimisation prediction using the UNAFold software version 3.4(47). In all cases N79 folding was robust (data not shown). We also introduced an RNA polymerase promoter immediately downstream of the 3’end of the genome in the plasmids encoding five of the N79-containing virus replicons to allow *in vitro* production of full length (gt2a HCV, HRV, HEV, CHIKV) or near full length (YFV) [-] RNA. Analysis of these RNAs showed that they experienced no more than a 2-fold difference in rates of cleavage, with cleavage ranging from 30% to 60% (Fig 5a). Equally importantly, these relatively small differences did not correlate with how effectively the N79 Rbz suppressed replication *in cellulo*, given that the antigenomic strand of HCV was the second least efficiently cleaved RNA species.

**Fig 5.**
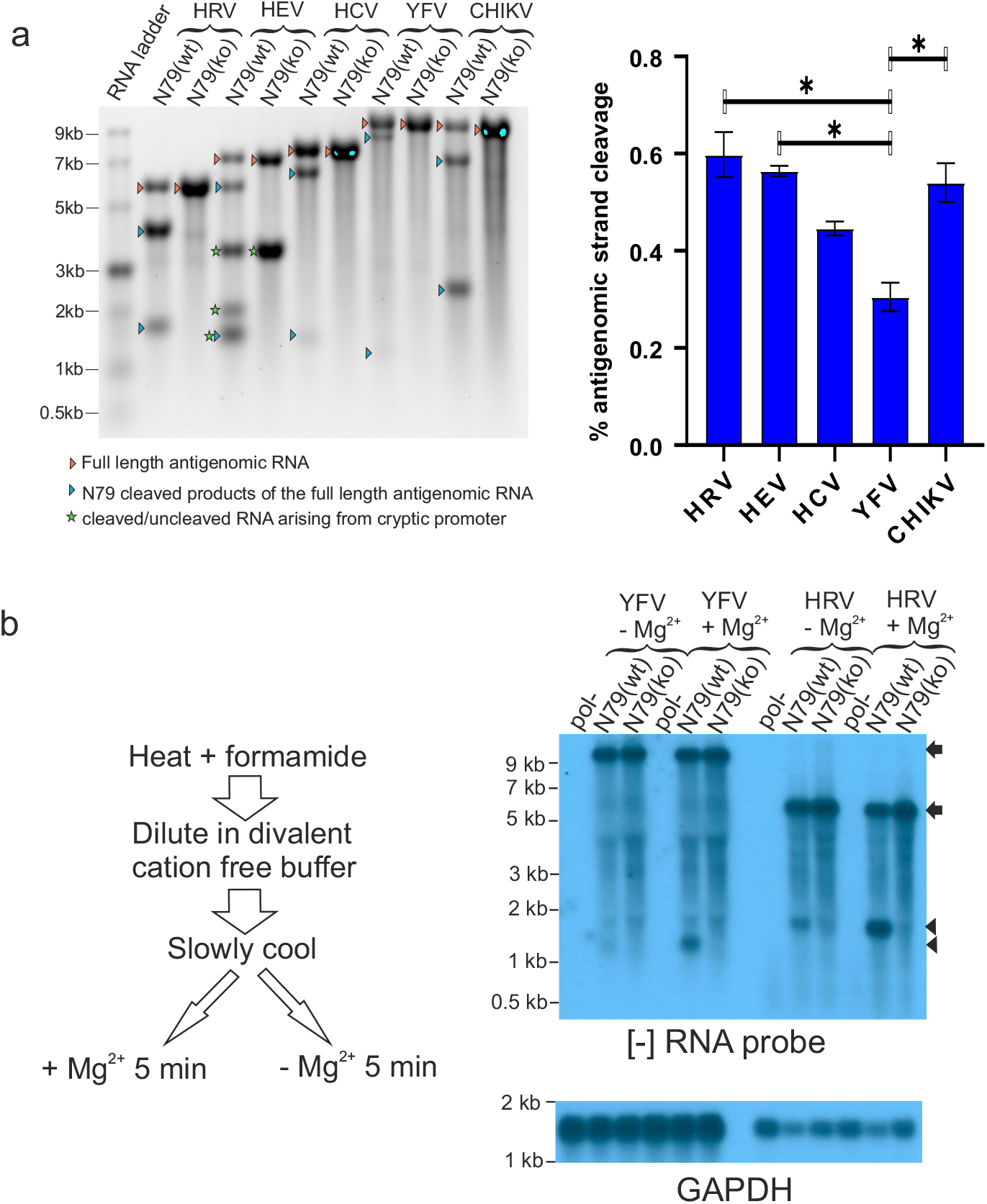
The N79 Rbz is not restricted in its ability to cleave the [-] RNA of positive strand RNA virus constructs when this RNA is in a single stranded state. (A) Single stranded [-] RNAs from various N79-containing replicons were generated by *in vitro* transcription and their cleavage was assessed by gel electrophoresis and image capture. Significant differences between experimental groups are highlighted (* p<0.05; one-way ANOVA; n=3). (B) RNA from cells transfected 24 hrs earlier with N79-containing YFV or HRV replicons was subject to treatment designed to release the [-] RNA from its double stranded state and enable it to fold before Mg^2+^ was added to activate Rbz activity. Experimental groups lacking Mg^2+^ addition were included as controls. RNAs were subjected to Northern blot analysis. The arrows represent the position of full length transcripts and arrowheads the position of N79-cleaved product.

Another scenario that might have accounted for poor cleavage is mutation of the Rbz sequence such that activity is rapidly lost. To exclude this possibility, cells were transfected with N79-containing HRV and YFV replicons and cellular RNA was recovered 48 hours later. Samples were heat denature to melt all dsRNA, then diluted and cooled to promote RNA folding. Half of each sample was supplemented with 1mM Mg^2+^ to enable Rbz catalysis while the other half was not. Subsequent Northern blot analysis using a probe complementary to the [-] RNA was used to assess cleavage of this RNA (Fig 5b). In the absence of Mg^2+^ supplementation, both YFV N79(wt) and N79(ko) transcripts appeared uncleaved and were equally abundant. The relative abundance of full length HRV [-] RNA N79(wt) and N79(ko) transcripts was also broadly similar, although unlike YFV a low level of cleavage of the N79(wt) transcript was also observed, consistent with the transient inhibition of replication observed at early time points in the replication assay. Importantly, the addition of Mg^2+^ resulted in an increase in cleavage of the HRV N79(wt) [-] transcript and cleavage of a proportion of the YFV N79(wt) [-] transcript not seen for the equivalent RNAs carrying the N79(ko) Rbz. We conclude a functional Rbz sequence is maintained in the [-] RNA strand of HRV and YFV during the time that genome amplification is occurring, but its activity is suppressed by dsRNA formation.

### Duplex formation blocks Rbz-mediated cleavage of NS5B synthesized products in vitro

The extent that Rbz induced cleavage of the [-] RNA of HCV was much greater than most other constructs tested. To investigate whether RNA synthesized by the HCV viral polymerase *in vitro* was equally prone to Rbz cleavage and thus able to sample secondary structure, we performed a series of RNA polymerase reactions. HCV NS5B RNA-dependent RNA polymerase reactions were set up to produce a 445 nt RNA transcript containing either an active (wt) or inactive N79 (ko) Rbz. Parallel control T7 DNA-dependent RNA polymerase reactions were set up to produce these same two transcripts as single-stranded RNA molecules. Analysis by denaturing polyacrylamide gel electrophoresis showed that about half of the N79(wt) RNA produced by T7 was cleaved into a ~400 and 90 nt product (Fig 6). This was specifically due to Rbz activity as the N79(ko) transcript produced by T7 remained uncleaved. In contrast, no obvious difference was observed between RNA products generated in the NS5B reactions when comparing between N79(wt) and N79(ko) experimental groups; the RNAs present consisting of an approximately unit length (~500 nt) product identical in size to the uncleaved RNA produced in the T7 reactions, as well as several smaller RNAs mostly ranging in size between 300-400 nts and likely to represent internal initiation products. To assess the double stranded nature of the various polymerase reactions, all products were incubated with RNase A under high salt conditions. No degradation of the NS5B products was seen, indicating that they were duplexed with their template and therefore double-stranded. In contrast the T7-derived products were highly sensitive to RNase A treatment. Finally, to confirm that dsRNA formation was preventing Rbz cleavage, the surviving RdRp product from the RNase treatment was subjected to heat denaturation and renaturation. In the N79(wt) experimental group this resulted in a decrease in size of all RNAs by approximately ~100 nt as well as the appearance of an additional ~90 nt band. In contrast there was no change in appearance of the N79(ko) containing RNAs. Taken together these data show that in the absence of any other host or viral encoded activity, RNA produced by HCV NS5B is unable to even transiently access secondary structures that might be encoded within it.

**Fig 6.**
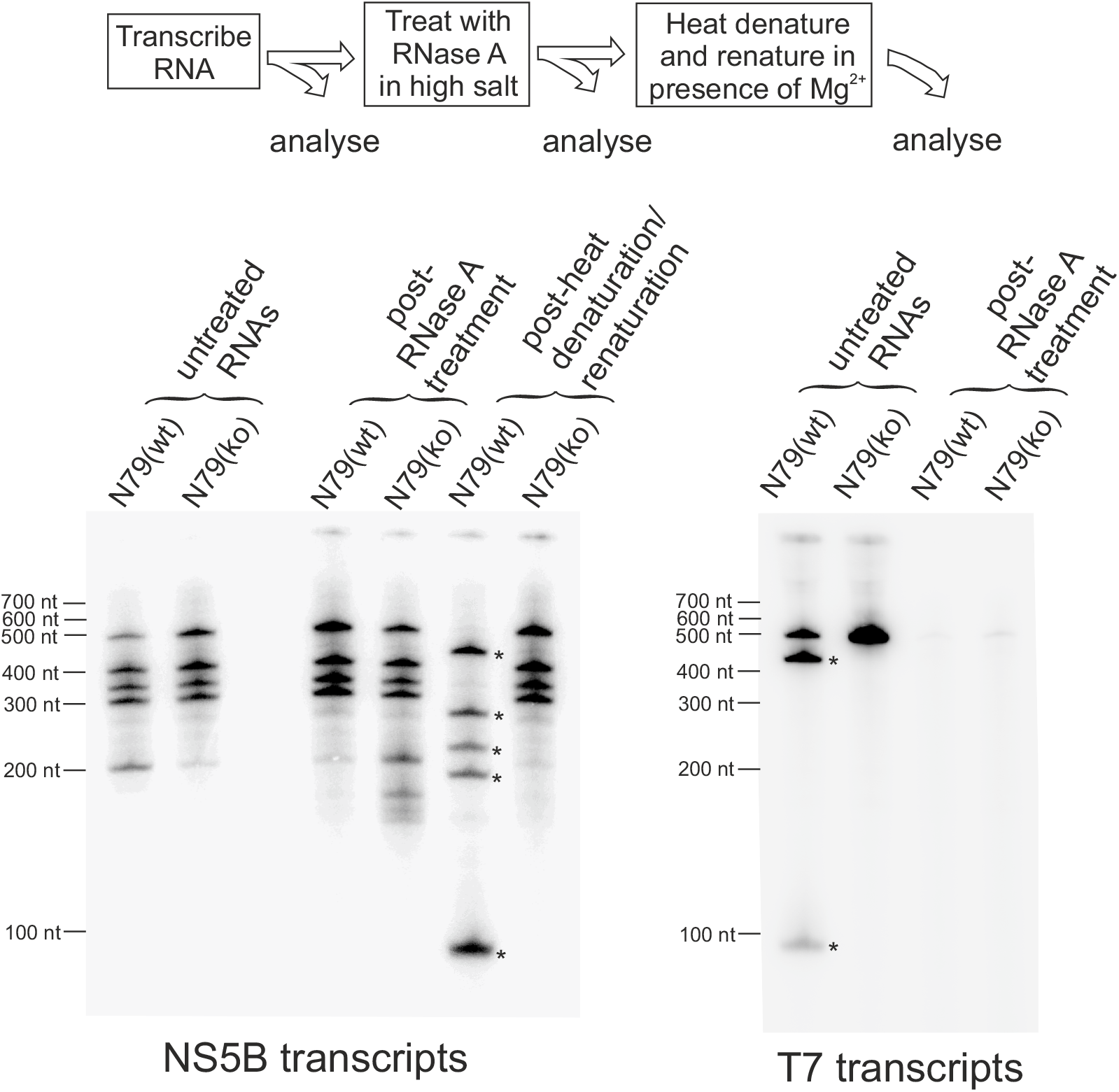
Rbz-containing RNAs are not cleaved when synthesised *in vitro* by the HCV RNA polymerase NS5B due to extensive base pairing with the template strand. A 490 nt RNA containing a reverse complemented active or inactive N79 Rbz was used as a template in an NS5B RdRp reaction. Products produced were [α-32P] labelled. A set of T7 reactions designed to synthesise identical RNA products from a DNA template were included as controls. All four RNAs were subjected to a series of treatments as detailed in the provided diagram before being run on a denaturing polyacrylamide gel. The high-salt RNaseA treatment step was used to selectively degrade single stranded RNAs. The heat denaturation/renaturation step was used to melt dsRNA and allow single stranded RNA folding. The astrices indicate RNAs arisen from Rbz cleavage. Results shown are representative of one of two experiments.

## DISCUSSION

Knowledge of the different structural states that RNA can adopt in the replication complexes of positive strand RNA viruses remains incomplete. For members of the *Leviviridae* family which infect prokaryotes, and for members of *Narnoviridae* which infect fungi and insects, the [+] and [-] RNAs are kept in a predominately single stranded state (30,48). In contrast, many other positive strand RNA viruses readily generate dsRNA when replicating (49,50). Indeed, this is such a ubiquitous property that plants and animals have evolved detection systems for dsRNA, recognition by which stimulates an array of antiviral responses (51,52). However, whether the viral [-] RNA is in a predominately single strand or double strand state when used as a template for [+] RNA synthesis has only been effectively answered for a few select viruses because of technical challenges addressing this question. One issue is that infection produces dsRNA through processes other than those involved in productive genome replication (53). Both this and the presence of exhausted replication complexes have the potential to generate dsRNA decoys containing [-] RNA. Another difficulty is that active replication complexes are protected by lipid membranes. Typically these need to be disrupted to gain experimental access to the RF and RI; a process which frustrates RNA structural analysis by promoting collapse of complementary RNAs into a double stranded state (31). Nonetheless, for some viruses there is good evidence to suggest that the RF and RI are double stranded (54–57). What is less clear is whether this is always the case. In this study, we aimed to address this gap in our knowledge by using Rbzs to assess RNA folding in the [-] RNA of various viruses. Our data show that the ability of a Rbz to cleave the [-] RNA of different viruses varies enormously. This observation is potentially consistent with the notion that the [-] RNA varies in the extent to which it is double stranded in different viral RCs. That said, the lack of Rbz of cleavage does not necessarily mean that a [-] RNA is in a double stranded state, as constraints may still be placed on its folding when it is single stranded. Indeed the limited cleavage of the HRV [-] by the N79 Rbz that we observed may reflect this, as the RI from poliovirus has previously been reported to be predominately single stranded (31). Constraints on folding of the [-] RNA could arise from it being bound by viral proteins, and explain why enterovirus such as poliovirus and HRV use structural elements at the 5’ of their [+] RNA as a promoter for [+] strand synthesis (58). Irrespective of the reasons, the fact that Rbz cleavage requires prior separation of dsRNA means that where extensive cleavage is observed, the [-] RNA either has to be predominately single stranded or be constantly fluctuating between a single and double stranded state.

The HCV RdRp, NS5B, produces dsRNA *in vitro* (59). Structural studies also suggest that newly synthesized products of NS5B leave the polymerase base-paired to their template (60). Our results confirm that nascent NS5B products produced *in vitro* fail to sample secondary structure as a result of this base pairing. Consequently, folding within the [-] RNA of HCV must depend on additional viral or host encoded activities such as the HCV helicase, NS3. Helicase activity is achieved by the protein binding to a stretch of single stranded RNA and migrating in a 3’-5’ direction, displacing any complementary RNA (61). Assuming NS3 is involved in opening up duplex structures in the RF and RI, one question that arises is how it accesses a region of single stranded RNA to enable binding. Perhaps access is gained at the ends of these RNAs, possibly as a result of duplex breathing or mediated by terminal transferase activity of NS5B providing a single stranded extension to one or other strands (62). If so, NS3 helicase processivity within the RC must be considerable to allow sampling of RNA structure as far in to the [-] strand as the position at which the Rbzs were placed. Another consideration is that once access is gained, any region of duplexed RNA opened up by the passage of NS3 still has the potential to reanneal once NS3 has moved on. As reannealing would reimpose a block on the [-] strand being able to fold, slowing or preventing it could also be important. It is possible that scaffolding activities provided by single stranded RNA binding proteins such as NS4B and NS5A (63,64), as well as NS3 working as a functional oligomeric array (65) play a role here. As RF and RI formation have yet to be formally demonstrated in HCV, it may even be possible that reannealing is prevented by physical separation of the [-] strand from its [+] counterpart. Indeed, *in situ* hybridization findings suggest that [-] RNA fails to co-localise with [+] RNA in HCV infected cells (66,67). However, it is difficult to reconcile such physical separation with the accepted model of HCV replication, that of an ER-derived membrane spherule harbouring both [+] and [-] RNA strands (68). Whatever mechanisms HCV and other viruses use to enable sampling of complex structures within their [-] RNA, a better understanding of these could provide novel therapeutic angles for treatment.

Production of a subgenomic RNA resulting from internal initiation within a full length [-] RNA is a feature common to the *Caliciviridae* (25), *Togaviridae* (69) and *Hepeviridae* virus families (70). In the case of the *Caliciviridae* family, genetic and biochemical evidence supports the involvement of a stem-loop found at the end of ORF1 as being critical in promoter function (25,45). For the *Togaviridae* family and other viruses within the so-called alphavirus super family, the role that RNA structure plays in subgenomic promoter activity is disputed (71–76). Subgenomic RNA production by members of the *Hepeviridae* family has been less extensively studied, in part because of historically less tractable cell culture systems. Nonetheless, the region within the viral genome harbouring the promoter has been identified (77), and [-] RNA conserved stem loop structures can be found at this location (78). Given the lack of impact N79 had on CHIKV, our study lends weight to the hypothesis that genomic and subgenomic promoters within the antigenomic strand of alphaviruses and *Togaviridae* more generally are recognised by their primary sequence. In contrast, the notable suppression of FCV by N79 and sTRSV Rbzs is consistent with the use of stem loop structures in the [-] RNA of the *Calicivirdae* family serving as a subgenomic promoters. Suppression of HEV replication by the N79 Rbz leaves open the possibility that structural elements harboured within the [-] RNA of *Hepeviridae* family members also serve as subgenomic promoters. However, it is notable that the extent to which HEV appears to sample RNA structure in this strand is far less than that of FCV.

Replication of West Nile Virus (WNV) has been suggested to depend on host cell proteins TIA-1 and TIAR binding to a 75 nt stem loop found within the 3’ end of the [-] strand (79). The helicase expressed by this and other flaviviruses also belongs to the same DEAD-box helicase superfamily as that expressed by HCV (80). Therefore, it was notable that introduction of the N79 Rbz, a RNA of similar size to the stem loop in the 3’ end of the [-] strand of YFV, had no impact on the replication of this virus. One of several reasons could account for this finding. Firstly, YFV and other flaviviruses may only be able to effectively sample RNA structure at the very ends of their [-] RNA because of reduced helicase processivity (81,82). Alternatively, reliance on TIA-1 and TIAR binding may not be a universal feature of flaviviruses. Finally it is possible that the predicted terminal stem-loop found at the end of the [-] RNA is not functional. Reverse genetics experiments looking at functional importance of putative stem loop functions at the terminal ends of the [-] RNA of positive strand viruses is fraught with difficulties. These complications arise because of the risk of off-target effects on structures within the genomic strand as well as the need to avoid synonymous changes in coding regions. While the lack of Rbz activity in the antigenomic strand cannot be used to disprove the existence of structural elements elsewhere in this same RNA, when Rbz cleavage does occurs it provides valuable supportive evidence that structural elements elsewhere are likely to form.

Our observations of Rbz activity within the antigenomic strand of HCV is consistent with the virus relying on a large structured promoter at the 3’ end of the [-] RNA. Even so it was noticeable how much more active Rbzs were in the HCV constructs compared to many of the other viral constructs. This begs the question, why does HCV expend what is likely to be a considerable amount of resources and energy driving duplex separation within its replication complex when it does not produce subgenomic RNAs. Indeed, it seems strange that any virus would evolve a large structured promoter within its [-] RNA unless there was already pre-existing evolutionary pressure driving stand separation. Perhaps by keeping its [+] and [-] strands in a more separated state, HCV reduces the chances of it being detected by dsRNA antiviral sensors. While the replication complex membranes protect the RF and RI from antiviral sensors in the cytosol, turnover/destruction of these organelles does expose them to endosomal TLR3, something that HCV already seeks to minimize by promoting RC secretion (83). By being single stranded when introduced into the endosome, complementary RNAs could be degraded before they form a more RNase-resistant double stranded state and therefore avoid TLR3 recognition. It is also possible that the protection afforded by the RC against dsRNA sensors is not absolute (4). Under such circumstances, having [-] and [+] RNAs in a more single stranded state would certainly be beneficial. Although HCV and other hepaciviruses are unusual amongst positive strand RNA viruses in their ability to establish a chronic infections with active viral replication in their immunocompetent hosts, they are not alone. Indeed, FCV achieves this (84), as does a number of other positive strand RNA viruses including members of the norovirus family (85), pegiviruses (86,87) and pestiviruses (88). In the future it would be interesting to determine the extent to which Rbzs are active in the [-] RNA of a broader range of these viruses, as well as examine whether changes to RNA structural sampling within this strand might impact on host dsRNA sensing.

## Supporting information

pictoral abstract

## DATA AVAILABILITY

Table S1 and Fig S1 are provided as supplementary files. Raw luciferase data as well as original .jpg, .tiff and .mp4 files produced during this study are available at https://doi.org/10.5281/zenodo.5873054

## FUNDING

This work was supported by the Medical Research Council [G0701215 to CJM; MR/S007229/1 to M.R.H; MR/S001026/1 to M.H.] and the Royal Society [RGS\R2\202376 to M.R.H.].

## ACKNOWLEDGEMENT

We would like to thank Professors A. Firth and D. Rowlands for helpful discussions. We are grateful to Prof T. Wakita, Prof R. Bartenschlager, Prof K. Green, Prof Andres Merits, Dr T. Tuthill, Dr P. Bredenbeek and Dr P. Farci for the gift of HCV gt2a, HCV gt1b, FCV, CHIKV, HRV, YFV and HEV constructs.

## CONFLICT OF INTEREST

None declared.

## Notes

### Competing Interest Statement

The authors have declared no competing interest.

