## Supplementary figure 1 for "Positive strand RNA viruses differ in the constraints they place on the folding of their negative strand"

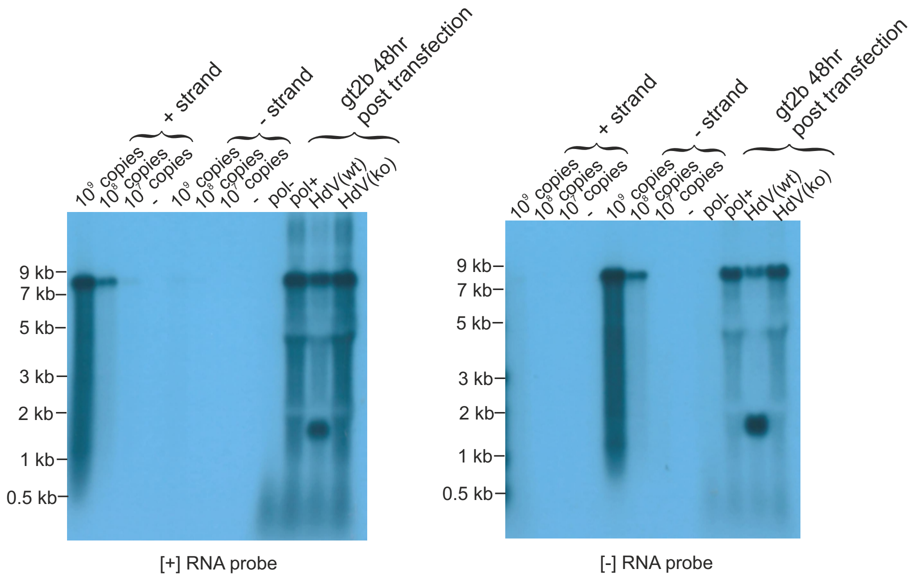


Figure S1. Strand specificity of sense and antisense probes directed at the Renilla luciferase coding region, present within all HCV, YFV and HRV replicon constructs used in this study. [+] and [-] strand RNA transcripts represent in vitro transcribed HCVgt2aN79(ko) and antisense HCVgt2aN79(ko) RNAs. Also present on the blot are cellular RNAs taken from cells 48 hours after transfection with various HCV gt2b constructs.
