## Supplementary table 1 for "Positive strand RNA viruses differ in the constraints they place on the folding of their negative strand"

### TABLE S1

| Primer number | Primer name | Sequence (5'-3') |
|---|---|---|
| 1 | Ren_HdV(fwd) | CAAAGCGGCCGTCCCATTCGCCATTAC |
| 2 | HdV_2A(rev) | GCCAGCAAGCGGCCGGCATGGTCCCAGCC |
| 3 | T7 | TAATACGACTCACTATAG |
| 4 | Ren_HdV(rev) | CGAATGGGACGGCCGCTTTGTTCATTTTTGAG |
| 5 | HdV_2A(fwd) | CATGCCGGCCGCTTGCTGGCAATCCACC |
| 6 | DVR4714-4694 | GTATGAGTGTCGTACAGC |
| 7 | HdVko(fwd) | GACGGTCCGCTGGGGATGTTGCCCAGCCGGCGCC |
| 8 | HdVko(rev) | CAACATCCCCAGCGGACCGTCCCCTCGGTAATG |
| 9 | JFH92-110 | GTATGAGTGTCGTACAGC |
| 10 | TSRV5’revflank | GCCGCTTTGTTCATTTTTGAG |
| 11 | TSRV3’fwdflank | TTGCTGGCAATCCACCC |
| 12 | DVR3511-3493 | GTTCTGTCTCTGCCGGTC |
| 13 | TSRVko(fwd) | GGACTCAAGTGACCGGAAAGCACATCC |
| 14 | TSRVko(rev) | CTTTCCGGTCACTTGAGTCCGTGAGGACGAAAC |
| 15 | N79ko(fwd) | CGAAGCGCGTCTCGTCCTATTTGGGACTC |
| 16 | N79ko(rev) | CAAATAGGACGAGACGCGCTTCGGTGCGTCC |
| 17 | DVR6677-6694 | CAACCTGAAGATCCCCTG |
| 18 | NS5ADVR_N79(rev) | CACTCATAGAGCAACAGACGGTAGTGTC |
| 19 | NS5ADVR_N79(fwd) | CGTCTGTTGCTCTATGAGTGGGGGCGGCGTGGATAGCAG |
| 20 | N79_NS5B(rev) | GTGGTATCATCCCCACCCCCACTGAGATGCA |
| 21 | N79_NS5B(fwd) | GTGGGGGTGGGGATGATACCACAGTTTGTTGTTC |
| 22 | DVR8867-8850 | AACTGTTTCCCAGGCAGC |
| 23 | AscI-Renilla(fwd) | CGCTGGGCGCGCCATGACTTCGAAAGTTTATGATC |
| 24 | VP1N79(rev) | CCAACAGTTGTTAGATTTGTCCCACCCCCACTGAGATGC |
| 25 | VP1N79(fwd) | GGACAAATCTAACAACTGTTGGGCCTAGTGACATGTATGTG |
| 26 | HRVA16(4053-4033) | TTGAGTTTCTTAAGTTCACTC |
| 27 | NotI-T7-5UTRYFV | GCGGCCGCTAATACGACTCACTATAGAGTAAATCCTGTGTGCTAATTG |
| 28 | YFVcapsid-Renilla(rev) | CGAAGTCATGCGAACTCCTCGTCGTACC |
| 29 | FMDV2A-YFVenv(fwd) | CCCCGGTCCGGTTGGCATCAACACAAGAAAC |
| 30 | NS1internal(rev) | TTCCGTTCACCGCTGCACC |
| 31 | YFVcapsid-Renilla(fwd) | GAGGAGTTCGCATGACTTCGAAAGTTTATGA |
| 32 | FMDV2A-YFVenv(rev) | GTTGATGCCAACCGGACCGGGGTTGGACTC |
| 33 | AvrII-deltaRen_N79(fwd) | CAGCTACCTAGGCCACCATGGAGCGAGTTCTCAAAAATGAAC |
| 34 | FF_FMDV2A(rev) | GGCGCTGGGCCCTTCTTAATGTTTTTGGCATCTTCCATCGGACCGGGGTTGGACTCG |
| 35 | HEV_nLuc_Fwd | CGCGACCATGAACTCCTTCTCCACAAGCGCC |
| 36 | HEV_nLuc_Rvs | GGAGTTCATGGTCGCGAACCCATGGG |
| 37 | HEV_2746_2769 | GGGGACGAGTTGTACCTTCC |
| 38 | HEV_5108_5089 | CATGGGCGCAGCAAAAGAC |
| 39 | EcoRI_extN79(fwd) | TAATAGGAATTCTGAGCGAGTTCTCAAAAATG |
| 40 | EcoRI_extN79(rev) | GTTATTGAATTCTCTGGCTTCAGTTGGGTG |
| 41 | HEV_RdRP_GNN_fwd | AAAGGTAATAACTCGATAGTGCTTTG |
| 42 | HEV_3’UTR_rvs | CAGGGAGCGCGAAACGCAG |
| 43 | HEV_RdRp_GNN_rvs | TATCGAGTTATTACCTTTAAAGGCAGC |
| 44 | HEV_3453_3473 | ACAGGGCGCTACCTACACAG |
| 45 | NotI_Ren(fwd) | ACAATTGGCGGCCGCTATCATAACCATGACTTCGAAAGTTTATGATCC |
| 46 | gt1bNS3_2A(rev) | GTAATAGGCGCCGGACCGGGGTTGGACTCGAC |
| 47 | 2A_gt1bNS3(fwd) | CCCCGGTCCGGCGCCTATTACGGCCTACTC |
| 48 | gt1b3flank | CTCGGTGGAGGGCATCTC |
| 49 | FCV (5247-5264) | CGCGGTCTGACCAGATCG |
| 50 | FCV MCS(rev) | TGCAGTCCGGAGGTGGTACCGTTTAAACACCAGTGCATTGGAGGGACAAC |
| 51 | FCV MCS(fwd) | GGTACCACCTCCGGACTGCAGAATGAGGCTGGAAAGATATTCC |
| 52 | FCV (6247-6228) | CTGTAACAATACACCCAGAG |
| 53 | XbaI_MfeI_5UTR(fwd) | TTAAGGTCTAGACAATTGACCTGCCCCTAATAGGGGCGACACT |
| 54 | UTR/FMDV fusion(1) | CTGAAGCCAGATGGTGCACGGTCTACGAGAC |
| 55 | UTR/FMDV fusion(2) | CCGTGCACCATCTGGCTTCAGTTGGGTG |
| 56 | T7_inner_Ren(fwd) | GAATTCTAATACGACTCACTATAGGCACCTGATGAAATGGG |
| 57 | 5’SGR | GCGACACGGAAATGTTGAATACTC |
| 58 | Xba_innerRen(fwd) | TTAAGGTCTAGAGCACCTGATGAAATGGG |

| | | |
|---|---|---|
| 59 | Renprobeouter(fwd) | CGAAAGGCCTTGTGGTACTGCCTGATAG |
| 60 | Renprobeouter(rev) | TTTTCTGTTTGTGTCTGGCTTCAGTTGG |
| 61 | Renprobeinner(fwd) | CGGGAGGTCTCGTAGAC |
| 62 | Renprobeinner(rev) | TAATGGCGAATGGGACG |
| **Ultra oligos** | | **Sequence (5'-3')** |
| N79 | | CTCAAAAATGAACAAAGCGGCGGGGGCGGCGTGGATAGCAGTGGAAT<br>CCAGGACGCACCGAAGCGCGTTTCGTCCTATTTGGGACTCATCAGCTGG<br>ATGTACCTGCATCTCAGTGGGGGTGGGTTGCTGGCAATCCACCCAAC |
| TSRV | | CTCAAAAATGAACAAAGCGGCTCTTCTTCTTCCTGTTTCGTCCTCACGGA<br>CTCATCAGACCGGAAAGCACATCCGGTGACAGGGTCTTCTTCTTTGCTG<br>GCAATCCACCCAAC |
| **Gblocks** | | **Sequence (5'-3')** |
| Nanoluc_antisense_sTSRV | | GTCCCTCCAATGCACTGGTGTTTAAACGGTACCATGGTCTTCACACTCG<br>AAGATTTCGTTGGGGACTGGCGACAGACAGCCGGCTACAACCTGGACC<br>AAGTCCTTGAACAGGGAGGTGTGTCCAGTTTGTTTCAGAATCTCGGGGT<br>GTCCGTAACTCCGATCCAAAGGATTGTCCTGAGCGGTGAAAATGGGCT<br>GAAGATCGACATCCATGTCATCATCCCGTATGAAGGTCTGAGCGGCGA<br>CCAAATGGGCCAGATCGAAAAAATTTTTAAGGTGGTGTACCCTGTGGA<br>TGATCATCACTTTAAGGTGATCCTGCACTATGGCACACTGGTAATCGAC<br>GGGGTTACGCCGAACATGATCGACTATTTCGGACGGCCGTATGAAGGC<br>ATCGCCGTGTTCGACGGCAAAAAGATCACTGTAACAGGGACCCTGTGG<br>AACGGCAACAAAATTATCGACGAGCGCCTGATCAACCCCGACGGCTCC<br>CTGCTGTTCCGAGTAACCATCAACGGAGTGACCGGCTGGCGGCTGTGC<br>GAACGCATTCTGGCGAGCGGCTCTTCTTCTTCCTGTTTCGTCCTCACGGA<br>CTCATCAGACCGGAAAGCACATCCGGTGACAGGGTCTTCTTCTCTGCAG<br>AATGAGGCTGGAAAGATATTCCAGC |
| Nanoluc_antisense_N79 | | GTCCCTCCAATGCACTGGTGTTTAAACGGTACCATGGTCTTCACACTCG<br>AAGATTTCGTTGGGGACTGGCGACAGACAGCCGGCTACAACCTGGACC<br>AAGTCCTTGAACAGGGAGGTGTGTCCAGTTTGTTTCAGAATCTCGGGGT<br>GTCCGTAACTCCGATCCAAAGGATTGTCCTGAGCGGTGAAAATGGGCT<br>GAAGATCGACATCCATGTCATCATCCCGTATGAAGGTCTGAGCGGCGA<br>CCAAATGGGCCAGATCGAAAAAATTTTTAAGGTGGTGTACCCTGTGGA<br>TGATCATCACTTTAAGGTGATCCTGCACTATGGCACACTGGTAATCGAC<br>GGGGTTACGCCGAACATGATCGACTATTTCGGACGGCCGTATGAAGGC<br>ATCGCCGTGTTCGACGGCAAAAAGATCACTGTAACAGGGACCCTGTGG<br>AACGGCAACAAAATTATCGACGAGCGCCTGATCAACCCCGACGGCTCC<br>CTGCTGTTCCGAGTAACCATCAACGGAGTGACCGGCTGGCGGCTGTGC<br>GAACGCATTCTGGCGAGCGGCGGGGGCGGCGTGGATAGCAGTGGAAT<br>CCAGGACGCACCGAAGCGCGTTTCGTCCTATTTGGGACTCATCAGCTGG<br>ATGTACCTGCATCTCAGTGGGGGTGGGCTGCAGAATGAGGCTGGAAA<br>GATATTCCAGC |
