## Supplementary material for "Positive strand RNA viruses differ in the constraints they place on the folding of their negative strand": pictoral abstract

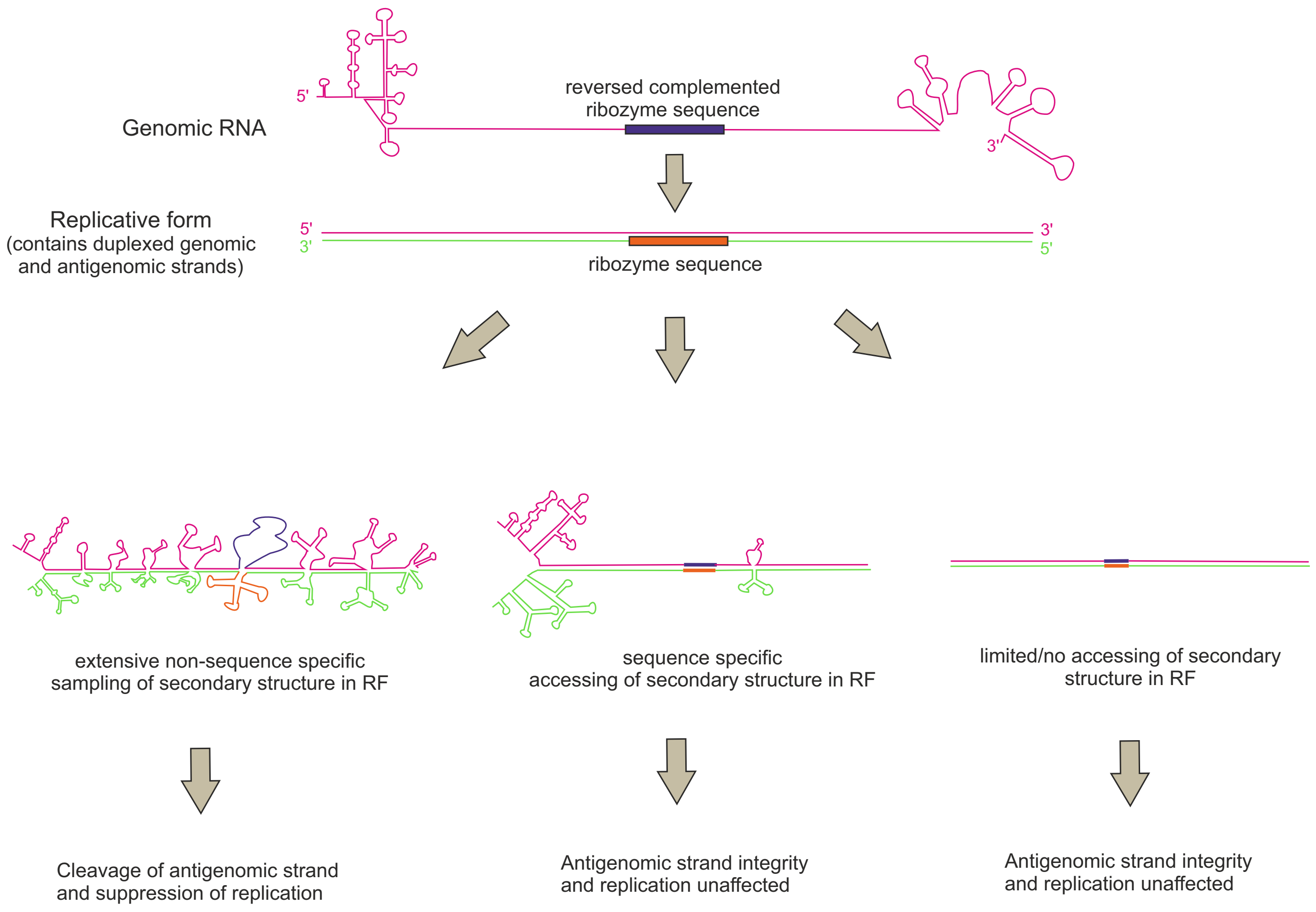
